# Ribosome fingerprinting with a solid-state nanopore

**DOI:** 10.1101/2020.08.07.236406

**Authors:** Mukhil Raveendran, Anna Rose Leach, Tayah Hopes, Julie L. Aspden, Paolo Actis

## Abstract

Nanopores hold great potential for the analysis of complex biological molecules at the single entity level. One particularly interesting macromolecular machine is the ribosome, responsible for translating mRNA into proteins. In this study, we use a solid-state nanopore to fingerprint 80S ribosomes and polysomes from a human neuronal cell line and, *Drosophila melanogaster* cultured cells and ovaries. Specifically, we show that the peak amplitude and dwell time characteristics of 80S ribosomes are distinct from polysomes and can be used to discriminate ribosomes from polysomes in mixed samples. Moreover, we are able to distinguish large polysomes, containing more than 7 ribosomes, from those containing 2-3 ribosomes, and demonstrate a correlation between polysome size and peak amplitude. This study highlights the application of solid-state nanopores as a rapid analytical tool for the detection and characterization of ribosomal complexes.

## Introduction

The analysis of individual biomolecular entities with nanopores is quickly emerging as a powerful bio-analytical tool. The technique is based on the principle of resistive pulse sensing^[1,2]^ wherein the temporary disruption in the measured ion current resulting from the passage of biomolecules through a nanopore is employed to study the size and conformation of the biomolecule. Over the years, nanopore sensing has been applied for numerous biomolecular analyses including DNA translocation studies, protein detection and nanopore-based nucleic acid sequencing^[3–5]^.

The single molecule sensitivity of the technique has also gained increasing attention in the field of RNA biology^[6]^ and in particular, biological nanopores have been widely used as RNA sequencing tools for molecular and clinical studies^[7–9]^. Besides sequencing, nanopores with their ability to distinguish small differences in charge and size of the translocating molecules are also excellent detection systems for studying the structure and conformation of RNA molecules. Accordingly, nanopores have been used for exploring RNA translocation dynamics^[10]^, tRNA translocation kinetics^[11]^ and to investigate the folding of RNA pseudoknot structures^[12]^. Nanopores have also been previously used to detect bacterial 50S ribosomal subunits and control their translocation^[13]^, and recently Rahman et al., reported the programmable delivery of 70S ribosomes with a nanopore integrated in an optofluidic chip^[14]^.

Despite these achievements, detection and analysis of individual ribosomes via nanopores has not yet been accomplished. Ribosomes are macromolecular machines comprising of RNAs and proteins that coordinate mRNA-guided peptide synthesis, a process known as translation. Eukaryotic ribosomes are composed of two subunits, the small 40S and the large 60S, which come together to form the 80S ribosomal complex during initiation of translation^[15,16]^. Each peptide synthesis event requires only one ribosome (monosome), however high levels of translation demand that multiple ribosomes bind an mRNA and synthesise peptides simultaneously. These multi-ribosome complexes are known as poly-ribosomes or polysomes^[17–19]^(Figure 1). Monosomes can represent efficient translation events but also spurious mRNA binding without generating a peptide, whereas the binding of multiple ribosomes to an mRNA is highly indicative of an active translation event^[18,20]^. Therefore, isolation and characterization of polysomes is important for gene expression and translational control studies^[21–24]^, specifically to investigate the levels of polysomes and monosomes, and their dynamics ^[20,25,26]^.

**Figure 1.**
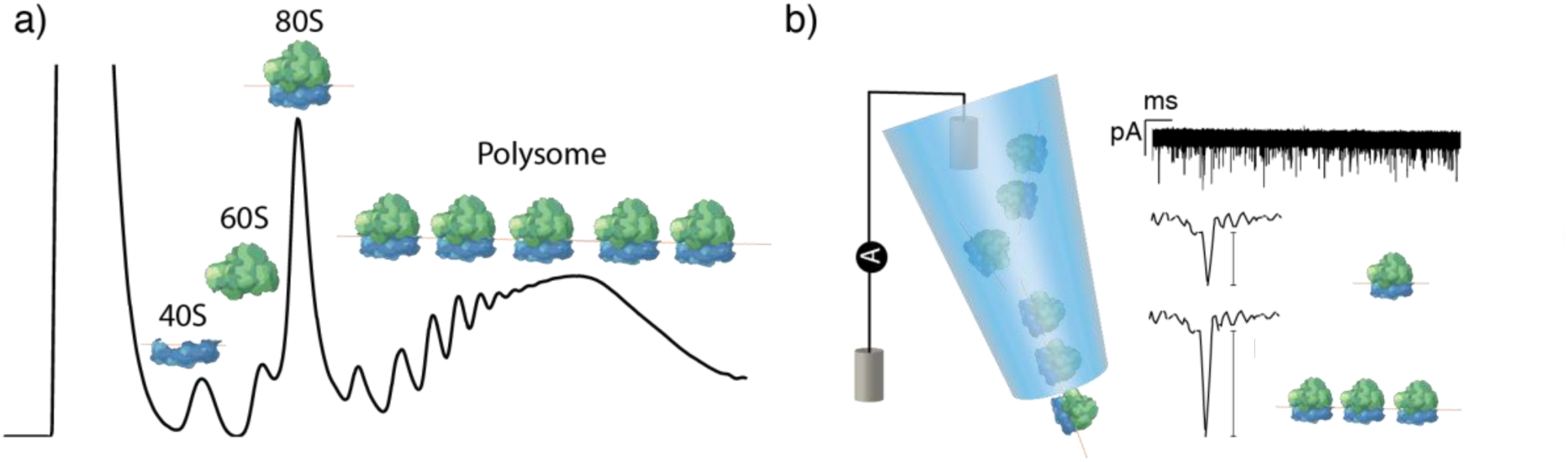
a) Schematic sucrose gradient UV trace from S2 cells indicating the types of ribosomal complex and how they are separated. b) Illustration of the nanopipette measurement setup and representative ion current signatures of monosomes and polysomes upon translocation.

In that regard, polysome profiling is a widely used approach to assess the global translational status of cells and tissues by separating large polysomes from small polysomes, single ribosomes and smaller RNA-protein complexes^[21][27]^. Typically, this separation is achieved via sucrose density gradient ultracentrifugation, which involves fractionation of sucrose gradients to isolate mRNAs with respect to number of bound ribosomes^[17,28]^. Other techniques include affinity purification and ultra-high-pressure liquid chromatography^[29]^. Although sucrose gradient ultracentrifugation is a prevalent method for polysome isolation and analysis, it requires sufficiently large quantities for data retrieval, which can be problematic for tissues or primary cells where material is limited. Moreover, the technique involves time consuming centrifugation, which could also lead to loss of bound proteins^[27,30]^.

Here, we propose solid-state nanopores as a quick and efficient analytical tool for the detection and analysis of ribosomes at the single-entity level. In this study, we show that characterization of nanopore ion current peak enables the detection of individual ribosomes and polysomes from both human and *Drosophila melanogaster* cultured cell lines, as well as *D. melanogaster* ovaries. Furthermore, we demonstrate the successful fingerprinting of polysome samples, differentiating large polysomes (>7 ribosomes bound to single mRNA) from polysomes with fewer ribosomes in small sample volumes of 3-5 µl. We believe solid state nanopores could become a useful tool for ribosome profiling to complement data generated from mass spectrometry and cryo-EM^[31–33]^.

## Results and Discussion

### Detection and analysis of 80S ribosomes

We investigated whether nanopores can detect ribosomes and polysomes with single entity resolution. Nanopore translocation experiments were carried out using quartz nanopipettes of ∼60 nm pore diameter, as shown in the SEM image (Figure 2a) with a corresponding resistance of 135±10 MΩ measured in 0.1 M KCl. The nanopipettes were produced via laser pulling using a two-line pulling parameters (Methods and materials). This method of nanopore fabrication is advantageous as it involves a straightforward bench top fabrication that produces low noise nanopores with consistent and reproducible pore sizes ^[34]^ as indicated in the current-voltage plots (Figure 2b). The ribosomal complexes of interest were introduced inside the nanopipette in 0.1 M KCl alongside the working electrode and a reference electrode in contact with the electrolyte was placed in an external bath. We have also tested the addition of Mg^2+^ to the measurement buffer to stabilize the ribosomes but noticed no difference in the recorded data for a 2-minute trace so we used 0.1M KCl for all experiments presented in this work. Upon application of a positive voltage to the electrode placed inside the nanopipette, the ribosome complexes translocate from inside to the outside of the nanopipette, causing a temporary decrease in the otherwise steady the ion current trace (Figure 2c).

**Figure 2.**
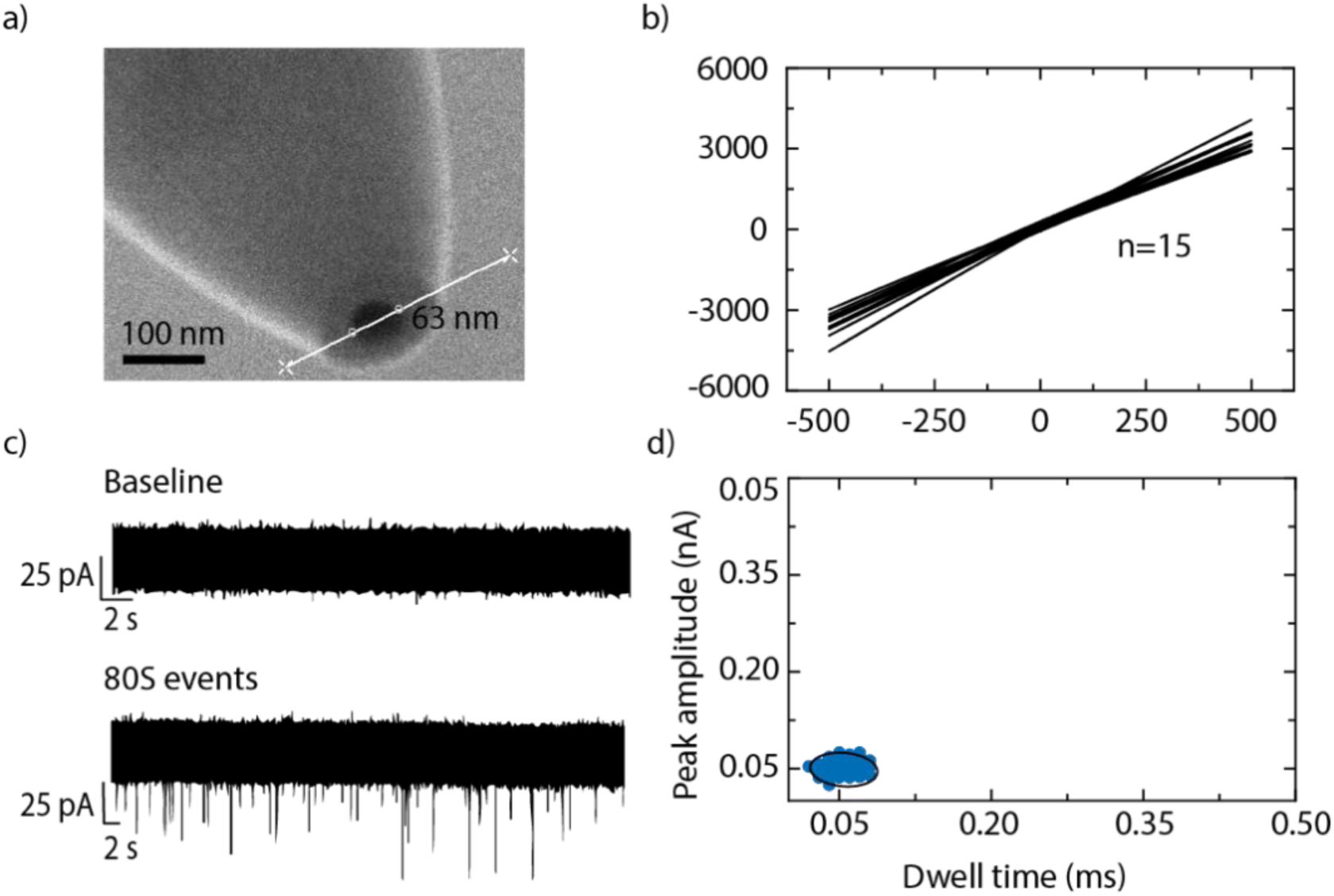
a) SEM micrograph of a nanopore at the tip of a nanopipette measuring approximately 63 nm in diameter b) IV characteristic of 15 nanopipettes fabricated using the same pulling parameter. c) Representative ion current trace in the absence (top panel) and presence (bottom panel) of ribosomes, each downward spike, called events, represent an individual 80S ribosome translocating out of the nanopore. d) Peak amplitude versus dwell time plot of 80S events of S2 cells indicating a tight cluster, the black line represents the 95% confidence ellipse.

80S ribosomes extracted from an embryonically derived *D. melanogaster* cell line (S2 cells) were examined to demonstrate the ability of nanopores to successfully detect individual ribosomes. These cells were chosen because it is straightforward to extract large quantities of ribosomes and levels of polysomes are relatively high (600 to 1400 µg/mL over the various gradients). Figure 2c shows a representative baseline ion current in the absence of 80S ribosomes (top trace) and an ion current trace in their presence (bottom trace), under a positive bias of 250mV. The downward spikes or events indicate individual ribosome translocations across the nanopore. No events were detected under an applied negative voltage indicating that the translocation of ribosomes is likely to be controlled by the electro-osmotic flow^[35]^. A typical ion current trace was recorded for two minutes.

Peak characteristic calculations for >100 such events presented a mean peak amplitude and dwell time of 40±5 pA and 0.07±0.01 ms respectively, as shown in the supplementary figure S1. These data indicate that 80S ribosomes translocate the nanopore causing an ion current blockade larger than 5σ from the average noise level (i.e. the false positive rate is ∼1 in 1,700,000) demonstrating the ability of solid-sate nanopores to detect 80S ribosomes. The scatter plot (Figure 2d) of the events with the individual peak amplitudes versus dwell time indicates that all of the observed events cluster together as a single population within a 95% confidence ellipse.

We also note that translocation of the small subunit of the ribosome, 40S ribosomal subunit, did not result in any detectable events (supplementary figure S2). This observation could be explained by considering the smaller size of the 40S subunit compared to the nanopore size used in this study or slightly different relative composition in terms of RNA and protein (40S: 53% rRNA and 47% protein; 80S: 61% rRNA and 39% protein).

### Differentiation of 80S monosomes from polysomes

Further, we demonstrate the ability of nanopores to differentiate 80S monosomes from small and large polysomes. We selected polysomes with varying number of ribosomes bound per mRNA, obtained from *D. melanogaster* S2 cells. The polysomes were separated into fractions via sucrose gradient ultracentrifugation (Methods and materials) and Figure 3a shows the separation of the polysomes into 6 fractions. The first four fractions contain approximately 2, 3, 4, 5 & 6 ribosomes referred to as p1, p2, p3, p4 respectively, the final two fractions are referred as p5 (∼7-11 ribosomes) and p6 (∼>11 ribosomes) respectively.

**Figure 3.**
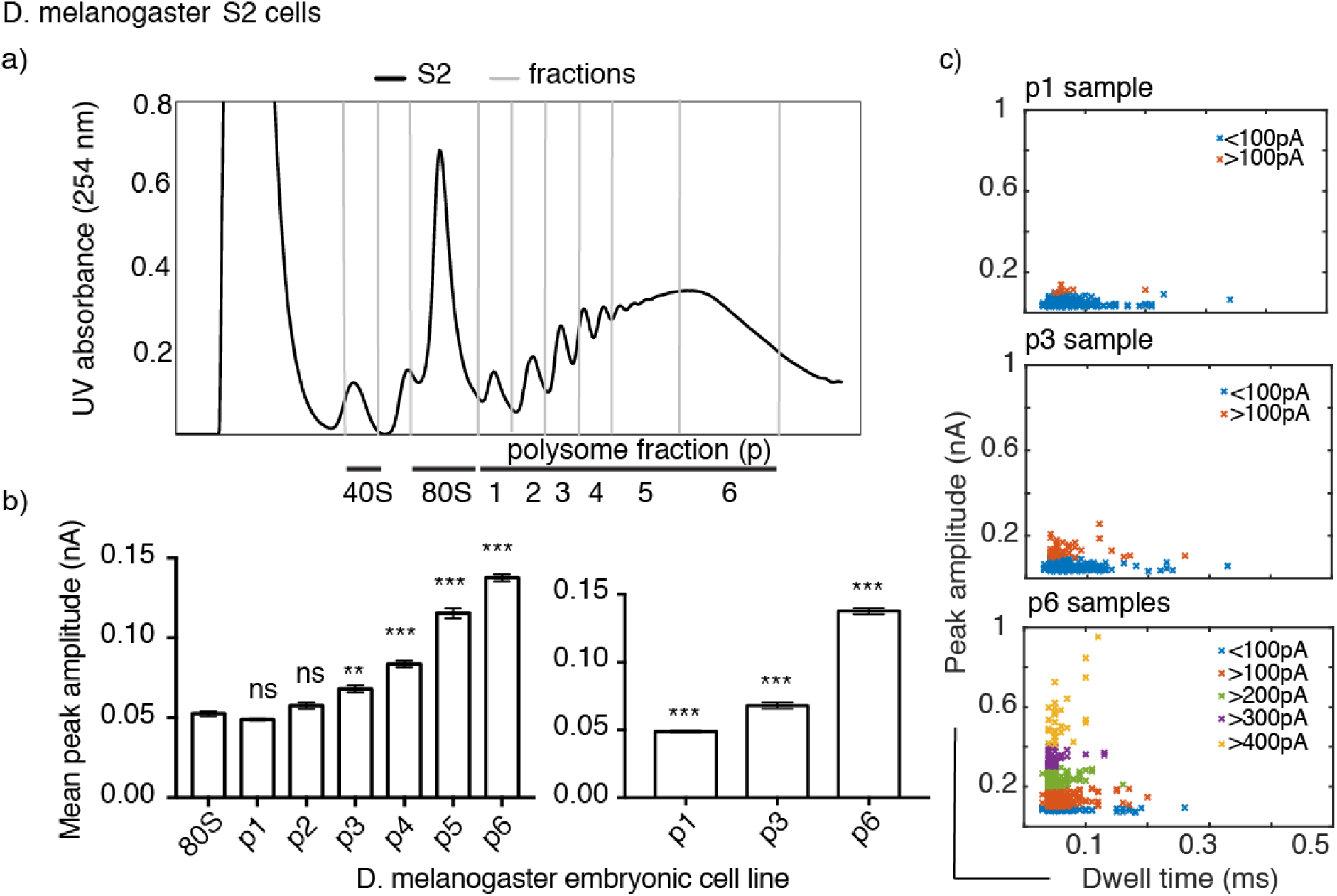
D. melanogaster S2 cell polysome analysis. a) UV (254nm) profile across sucrose gradient with ribosomal complexes separated based on their sedimentation profile. RNP (ribonucleoproteins and RNA) remain at the top of the gradient, followed by 40S subunits, 60S, 80S and polysomes with various numbers of ribosomes bound per mRNA. Up to 11 ribosomes per mRNA can been seen in this example, fractions were taken as indicated p1, p2, p3, p4, p5, and p6. b) Statistical Kruskal-Wallis test for each of the polysome samples against 80S shows that there is no significant difference (ns) in the peak amplitude for p1 and p2, whereas there is a significant difference with p value of 0.03 (*) for p3 and p<0.001 for p4, p5 and p6 indicated by ***. Similar tests performed for p1, p3 and p6 shows that there is a significant difference between these three polysome samples, with p<0.001 indicated by ***. The error bars indicate standard error of mean of each sample. c) Peak amplitude plotted against dwell time for at least 100 individual events for each sample, the events are colour coded to represent every 100 pA peak amplitude increase.

We analyzed the polysome samples with the nanopore setup and studied the resulting ion current events under the same conditions as before. Ion current event characterization of the different samples revealed an increasing mean peak amplitude in relation to the ribosome number per mRNA (supplementary Figure S3), with a significant difference (p<0.001) as shown in Figure 3b. This result can be explained by the increase in the magnitude of the ion current blockade caused by an increase in the volume occupied by the sample inside the nanopore. Supplementary figure S3 shows the peak amplitude histogram for the different polysome samples. The histograms indicate the occurrence of a second cluster of higher peak amplitudes, which is in correlation with the number of ribosomes present in the polysome. For example, p1 samples which contain mRNAs bound by ∼2 ribosomes exhibited a mean peak amplitude of 40±10 pA, whereas p3 bound by ∼4 ribosomes and p6 bound by ∼>11 ribosomes exhibited two mean peak amplitudes (p3 = 44±6 pA and 66±10 pA, p6 = 89±15 pA and 146±20 pA). Particularly, p6 samples exhibited very high peak amplitudes distributed over a wide range that were absent for samples with fewer ribosomes per mRNA.

In comparison, the dwell time analysis resulted in small differences between the different polysomes fractions with a varied distribution of events in relation to the ribosome number present in the sample (supplementary Figure S3 and S5). The long dwell time events and widespread peak amplitudes could be due to multiple polysomes translocating the nanopore at the same time or long polysomes translocating the nanopore in various conformations (circularized/linear). Individual histogram plots of peak amplitude and dwell time for all the polysome samples are provided in the supplementary Figure S3.

From the analysed samples, it is clear that the polysomes exhibit a higher mean peak amplitude than the 80S ribosomes. In fact, polysomes containing >7 ribosomes per mRNA, could be identified from those with fewer. Figure 3c depicts the scatter plots for polysomes p1, p3 and p6 demonstrating the occurrence of ion current events with a peak amplitude of >100 pA, >200 pA and >300 pA respectively. While the p1 sample has most of its ion current events within 100 pA, the p3 sample exhibits events in two groups, below and above 100 pA. This increase in peak amplitude is due to an increase in ribosome number (supplementary figure S4 and S5), and for p6 samples 80% of the events exhibit a peak amplitude of >100 pA as shown by the differently colored clusters in Figure 3C. The noted overlap in peak amplitude between the different polysome samples could stem from consecutive fractions containing populations with overlapping number of ribosomes per mRNA, additionally the varying RNA-protein complexes between mRNAs could also contribute to this effect. Nevertheless, the nanopores are sensitive enough to detect and differentiate 80S ribosomes from polysomes with 2-3 and >7 ribosomes per mRNA. The clear distinction between 80S ribosome samples and those containing polysomes could therefore be utilized as a quick screening technique.

Nanopore fingerprinting was also tested with lysates purified from *D. melanogaster* ovaries, wherein the concentration of polysomes (280 and 830 µg/mL) obtained was much lower than that in the cell lines. Cytoplasmic lysate from ovaries was subjected to polysome profiling and the isolated 80S ribosomes and polysomal complexes referred to as O80S and Op were then subjected to nanopore analysis (Figure 4a).

**Figure 4.**
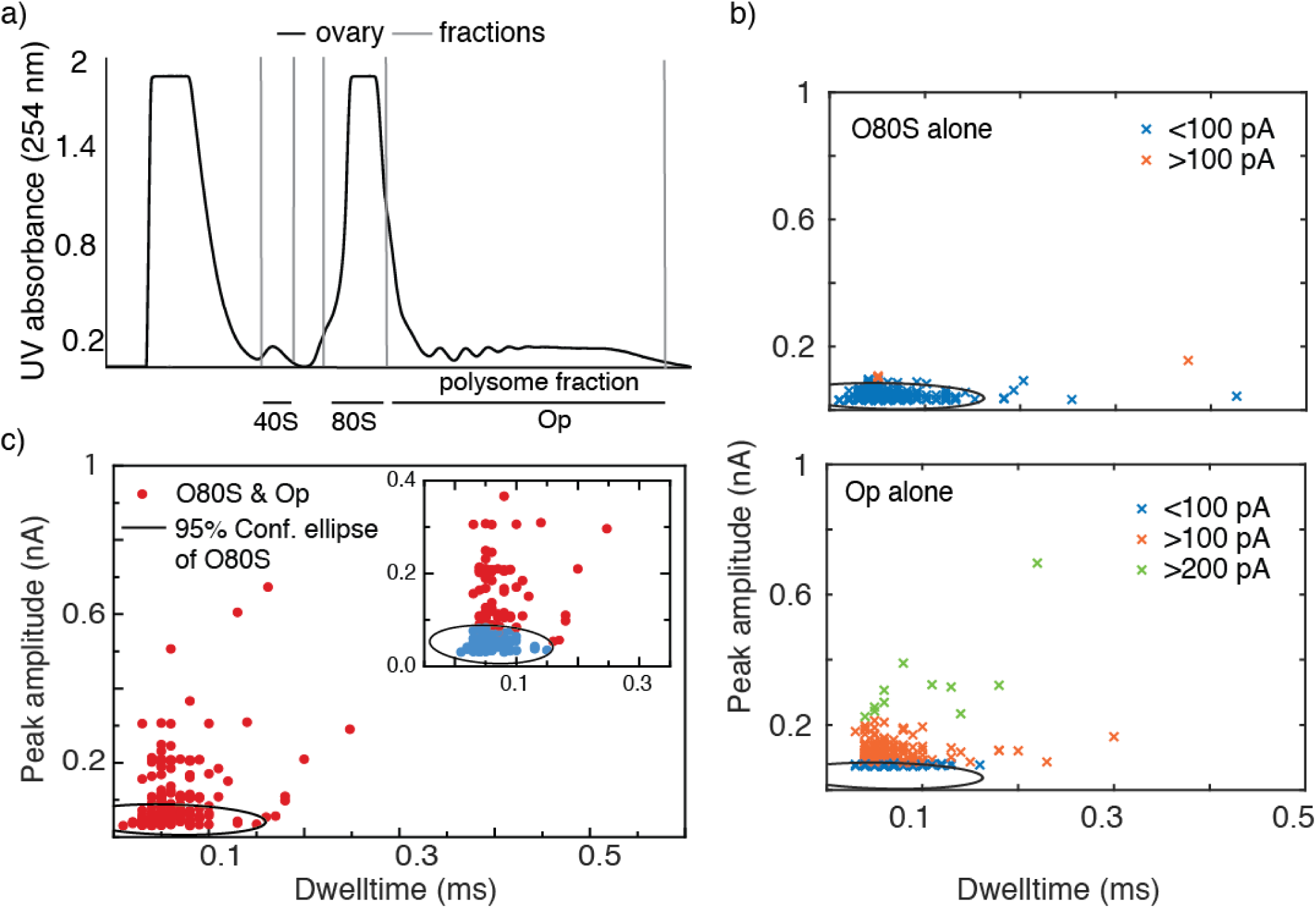
a) Sucrose gradient fractions of lysates purified from D. melanogaster ovaries separated into 40S, 80S and polysomes (p). b) Scatter plots for individual 80S and polysome samples obtained from D. melanogaster ovaries, indicating the clear difference in ion current peak characteristics between the two samples. The black circle indicates the 95% confidence ellipse fitted for 80S data and superimposed onto the polysome data. c) Peak amplitude plotted against dwell time for 80S and polysome mixture, the zoomed in inset indicates the events that fall within the 95% confidence ellipse represented in blue.

Similar nanopore translocation studies revealed that the O80S samples exhibit ion current events with a mean peak amplitude of 41±12 pA and dwell time of 0.05±0.01 ms, which closely match the data obtained for S2 cell 80S samples. As expected, the polysome samples, exhibited significantly different peak amplitudes of >100 pA (figure 4b), when compared to O80S, with a distribution similar to that of the S2 cell polysome samples (p<0.001, supplementary figure S9). Furthermore, these results were compared with a *D. melanogaster* ovary sample containing an 80S and polysome mixture to see if we could detect proportions of 80S and polysomes in a complex sample. Figure 4c shows the translocation events observed for the mixed fractionate indicating two slightly overlapping clusters, one below 100 pA and the other above 100 pA, which relates well with the two individual samples. As shown in the inset of Figure 4c, the 95% confidence ellipse fitted for the individual 80S samples was used as a boundary to identify the 80S events in the mixed sample. The results specify the potential of nanopores to accurately distinguish between single 80S ribosomes and polysomes in an unfractionated mixture. Similar studies for mixed 80S and polysome samples for S2 cells are provided in the supplementary Figure S6.

Having validated the nanopore analysis of ribosome extracted from a *D. melanogaster* cell line and from an ovary sample, we then tested the ability of the nanopore platform to analyze human ribosomes, extracted from SH-SY5Y human neuronal cell line. Here, in addition to the 80S units, polysome fractions with 2-5, 6-∼11, and ∼>11 ribosomes (referred to as hp1, hp2 and hp3) were purified. The analysis of the events showed similar peak amplitude and dwell time distinctions for 80S ribosomes and polysomes as measured for *D. melanogaster* S2 cells and ovaries. Figure 5b shows the discrete differences in the peak amplitude and dwell time scatter plots for the samples (80S ribosomes and polysome fractions), with 80S exhibiting 32±4 pA mean peak amplitude and 0.07±0.02 ms mean dwell time. Again for the polysomes, as observed for S2 cells, there is an increase in peak amplitude with respect to the ribosome number. While hp1 samples exhibited a mean peak amplitude of 53±12 pA, the samples with higher ribosome numbers exhibited two groups with mean of 56±10 pA and 91±20 pA for hp2, and 63±11 pA and 98±30 pA for hp3. However, all three polysome samples showed similar mean dwell times of 0.08±0.03 ms, 0.07±0.02 ms, 0.07±0.02 ms respectively, with the large polysomes (hp2 and hp3) exhibiting a wider distribution than the hp1 and 80S samples (Supplementary Figure S11).

**Figure 5.**
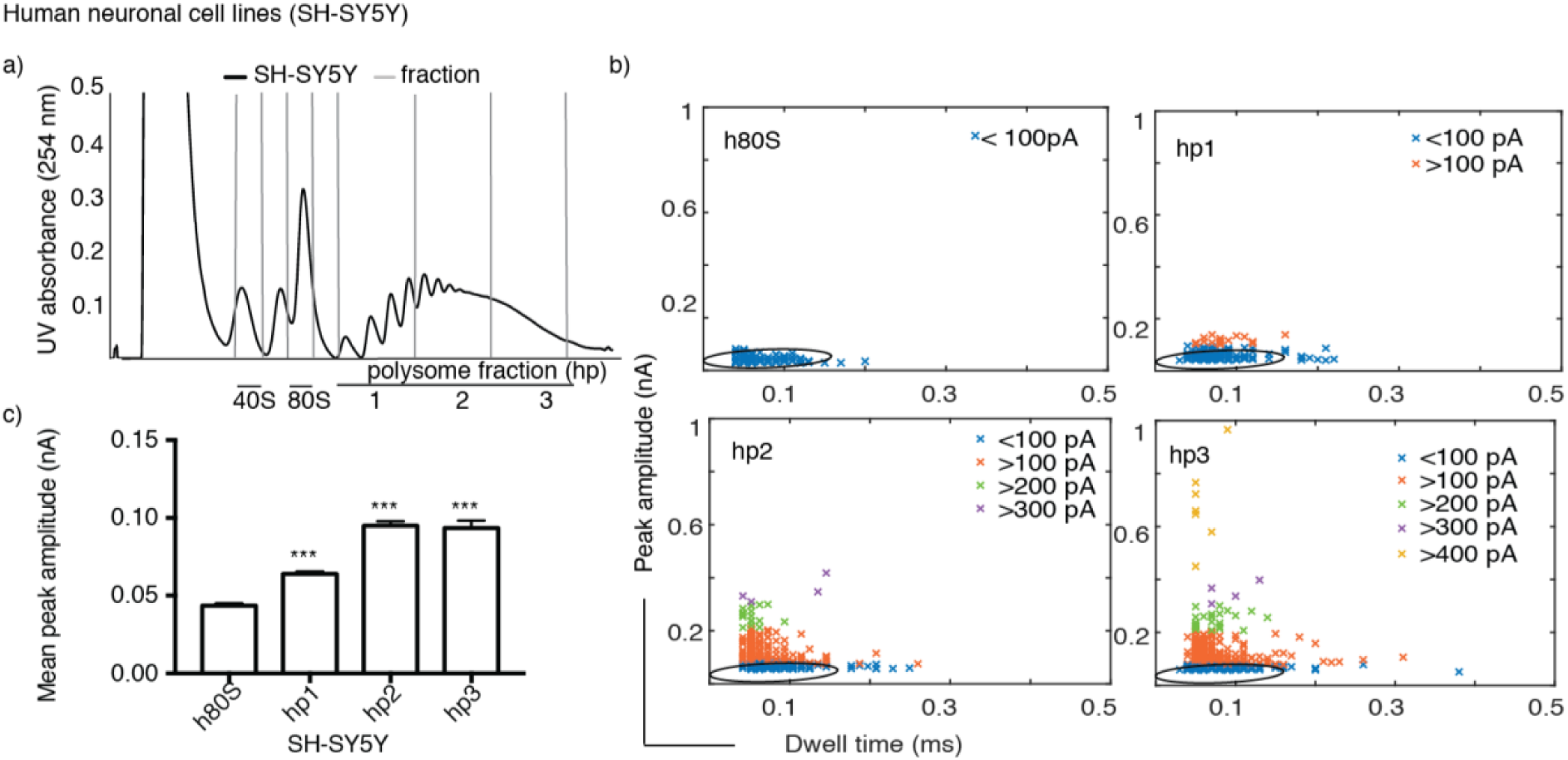
a) Sucrose gradient UV trace for human neuronal cell lines sample (SH-SY5Y), the polysomes are fractioned into hp1, hp2, and hp3. b) Scatter plots of 80S and polysomes (with 2-5, 6-∼11 and ∼>11 ribosomes), the graphs show a similar trend of peak amplitude increase with respect to increase in the number of ribosomes. The h80S data is fitted with a 95% confidence ellipse, which is then superimposed onto the polysome data. c) Mean peak amplitude data of SH-SY5Y polysomes compared with 80S samples exhibit a significant difference of p<0.001 represented by ***. Kruskal-Wallis test was performed for this data and the error bars indicate standard error of mean.

## Conclusions

In this study, we report the application of a solid-state nanopore for the analytical detection of ribosomes and polysomes. Characterization of these modulations allows us to detect single ribosomes and polysomes in very small sample volumes (3-5µl). We demonstrate that there is a significant difference in the peak amplitude between samples containing 80S ribosomes and those containing polysomes. Specifically, we provide evidence for the correlation between the number of ribosomes in a polysome and the resulting peak amplitude. These observations are consistent across samples derived from *D. melanogaster* S2 cells, *D. melanogaster* ovaries and SH-SY5Y human neuronal cells.

Thus, as well as being able to detect ribosomes and polysomes, these experiments reveal the ability of solid-state nanopores to differentiate between large polysomes (>7 ribosomes) from polysomes with a lower ribosome count, using their peak amplitude. Furthermore, using the same technique we generate characteristic fingerprints for single 80S ribosomes allowing us to distinguish between 80S and polysomes in mixed samples, indicating the robustness of the technique as an analytical tool.

Further extension of this study to achieve a greater separation between consecutive ribosome numbers and increased sensitivity of the nanopore to fully map the polysomes is foreseeable in the future, for example by taking advantage of the signal enhancement generated by macromolecular crowding^[36]^ or by careful selection of the electrolyte^[37]^. This could pave the way for the use of solid-state nanopores to distinguish individual polysome fractions as an alternative or complement to sucrose density gradient ultracentrifugation but with the advantage of doing so at the single entity level. The current study proves the potential of nanopores as a sensitive tool for identification of ribosomes and additionally to differentiate them from polysomes in a sample. We expect that with further improvements solid state nanopores can be a useful analytical tool, complementing cryo-EM and mass spectrometry for the structural analysis of ribosomes and ribonucleic particles.

## Methods and materials

### Cell culture

Semi-adherent S2 embryonic cells were maintained in Schneider’s medium containing L-glutamine (Sigma) supplemented with 10% FBS (Sigma), 100 U/mL penicillin, 100 µg/mL streptomycin, 25 µg/mL amphotericin B (GE Healthcare) and maintained at 26°C in non-vented, adherent flasks (Sarstedt). Human neuroblastoma SH-SY5Y cells, were cultured in Dulbecco’s Modified Eagle Medium (DMEM 4.5g/L Glucose with L-Glutamine) supplemented with 10% Fetal Bovine Serum (FBS) and 1% (v/v) Penicillin/Streptomycin at 37°C and 5% CO_2_.

### *D. melanogaster* husbandry

*D. melanogaster* wild type (Dahomey) were raised on standard sugar–yeast agar (SYA). Flies were kept at 25°C and 50% humidity with a 12:12 hr light:dark cycle in 6 oz Square Bottom Bottles (Flystuff). Semi-adherent S2 cells were maintained in Schneider’s medium containing L-glutamine (Sigma) supplemented with 10% FBS (Sigma), 100 U/mL penicillin, 100 µg/mL streptomycin, 25 µg/mL amphotericin B (GE Healthcare) and maintained at 26°C in non-vented, adherent flasks (Sarstedt).

### Ribosome purification and quantification

All stages were performed on ice or at 4°C wherever possible and all solutions were pre-chilled to 4°C. ∼300 pairs of ovaries were harvested from 3-6 day old females in 1X PBS (Lonza) with 1 mM DTT (Sigma) and 1 U/µL RNasin Plus (Promega) and flash frozen in liquid nitrogen. Ovaries were disrupted using RNase-free 1.5mL pestles (SLS) in lysis buffer (50 mM Tris-HCl pH 8 (Sigma), 150 mM NaCl, 10 mM MgCl_2_ (Fluka), 1% IGEPAL CA-630 (Sigma), 1 mM DTT, 100 µg/mL cycloheximide, 2 U/µL Turbo DNase (Thermo Fisher), 0.2 U/µL RNasin Plus, 1X EDTA-free protease inhibitor cocktail (Roche)). SH-SY5Y cells and S2 cells were treated with 100 µg/mL cycloheximide (Sigma) for 3 minutes before harvesting. Cells were pelleted at 800 x g for 8 minutes, washed in ice-cold 1X PBS supplemented with 100 µg/mL cycloheximide. Ovaries, SH-SY5Y cells and S2 cells were lysed in 500 µL lysis buffer for ≥30 mins with occasional agitation, then centrifuged for 5 minutes at 17,000 x g to remove nuclei.

Cytoplasmic lysates were loaded onto step-wise 18 – 60% sucrose gradients (50 mM Tris-HCl pH 8.0, 150 mM NaCl, 10 mM MgCl_2_, 100 µg/mL cycloheximide, 1 mM DTT, 1X EDTA-free protease inhibitor cocktail) and ultra-centrifuged in SW40Ti rotor (Beckman) for 3.5 h at 170,920 x g at 4°C. 0.5 mL fractions were collected using a Gradient Station (Biocomp) equipped with a fraction collector (Gilson) and Econo UV monitor (BioRad). Fractions were combined according to polysome peaks, diluted to 10% sucrose, concentrated using 30 kDa column (Amicon Ultra-4 or Ultra-15) and buffer exchanged (50 mM Tris-HCl pH 8, 150 mM NaCl, 10 mM MgCl_2_) at 4°C until final sucrose percentage of ≥0.1%. Samples were quantified using Qubit Protein Assay Kit, diluted 20-fold in filter sterilised nano pipette translocation buffer (0.1 M KCl) containing 10 mM MgCl_2_ then frozen using liquid nitrogen in single-use aliquots.

### Nanopipette fabrication

The nanopipettes with ∼60 nm diameters were fabricated from quartz glass capillaries of 0.5 mm inner diameter (QF100-50-7.5, World precision Instruments, UK) using a Sutter instrument model P-2000 laser puller. The pulling protocol comprised two separate lines with the parameters line 1: HEAT 775 FIL 4 VEL 30 DEL 170 PULL 120 and line 2: HEAT 900 FIL 3 VEL 20 DEL 175 PULL 180. Using these parameters, pulling highly consistent glass nanopipettes with pore sizes with variations of less than 10 nm was possible with pipettes pulled on different days. Ag/AgCl wires (0.25mm diameter, Sigma Aldrich, UK) were utilized as both the working and counter electrodes.

### Ion current measurements

For the translocation experiments nanopipettes fitted with the working electrode were filled with translocation buffer (0.1 M KCl) containing the ribosome samples at a final concentration of 20µg/ml. The nanopipette and a grounded reference electrode were immersed in a 0.1 M KCl solution completing the circuit. On application of a positive potential to the working electrode, ribosomes from inside the nanopipette translocate through the nanopipette pore into the electrolyte solution, resulting in a temporary blockage of the ion current. Ion current data were acquired using an Axon instruments-patch clamp system (Molecular devices, USA). Measurements were recorded using the Axopatch 700b amplifier, and the data were acquired at a rate of 100 kHz and low pass filtered at 20 kHz using Pclamp 10.6 software. Initial data analysis was carried out with a custom MATLAB script (provided by Prof Joshua Edel, Imperial College London, UK) and further data analysis was carried out using proFit 7 (QuanSoft, Switzerland).

## Acknowledgements

JA and PA are funded by the University of Leeds (University Academic Fellow scheme). This work was funded by Royal Society (RSG\R1\180102), BBSRC (BB/S007407/1), Wellcome Trust ISSF (105615/Z/14/Z) and White Rose University Consortium-Collaborative Grant. ARL acknowledges a School of Electronic and Electrical Engineering Scholarship from the University of Leeds. MR acknowledges the LARS Scholarship by the University of Leeds. We thank Prof Joshua Edel (Imperial College London) for sharing a custom MatLab script for single molecule analysis.

## Author Contribution

M.R. and A. R. L performed the nanopore experiments, performed the data analysis and wrote the first draft of the manuscript. T.H. quantified and purified the ribosomes. P.A. and J. L. A. designed and supervised the project. All authors discussed the experiments and approved the manuscript.

## Notes

The authors declare no competing financial interest

## Data availability

Data supporting this work can be accessed via the University of Leeds repository: TBD

## References

[1] Y. Song, J. Zhang, and D. Li, “Microfluidic and nanofluidic resistive pulse sensing: A review,” Micromachines, vol. 8, no. 7, pp. 1–19, 2017.

[2] H. Bayley and C. R. Martin, “Resistive-Pulse Sensing From Microbes to Molecules,” Chem. Rev., vol. 100, no. 7, pp. 2575–2594, 2000.

[3] W. Shi, A. K. Friedman, and L. A. Baker, “Nanopore Sensing,” Anal. Chemisry, vol. 89, pp. 157–188, 2017.

[4] C. Dekker, “Solid-state nanopores,” Nat. Nanotechnol., vol. 2, no. 4, pp. 209–215, 2007.

[5] T. Albrecht, “Single-Molecule Analysis with Solid State Nanopores,” Annu. Rev. Anal. Chem., vol. 12, pp. 371–387, 2019.

[6] R. Y. Henley, S. Carson, and M. Wanunu, “Studies of RNA Sequence and Sturcture using Nanopores,” Prog Mol Biol Transl Sci, vol. 139, pp. 73–99, 2016.

[7] N. Kono and K. Arakawa, “Nanopore sequencing: Review of potential applications in functional genomics,” Dev. Growth Differ., vol. 61, no. 5, pp. 316–326, 2019.

[8] A. Mueller, K. Fischer, R. Suluku, and T. Hoenen, “Sequencing of mRNA from whole blood using nanopore sequencing,” J. Vis. Exp., vol. 2019, no. 148, pp. 1–8, 2019.

[9] M. T. Parker et al., “Nanopore direct RNA sequencing maps the complexity of arabidopsis mRNA processing and m6A modification,” Elife, vol. 9, pp. 1–35, 2020.

[10] C. Shasha, R. Y. Henley, D. H. Stoloff, K. D. Rynearson, T. Hermann, and M. Wanunu, “Nanopore-Based Conformational Analysis of a Viral RNA Drug Target,” ACS Nano, vol. 8, no. 6, pp. 6425–6430, 2014.

[11] R. Y. Henley, B. A. Ashcroft, I. Farrell, B. S. Cooperman, S. M. Lindsay, and M. Wanunu, “Electrophoretic Deformation of Individual Transfer RNA Molecules Reveals Their Identity,” Nano Lett., vol. 16, no. 1, pp. 138–144, 2016.

[12] X. Zhang et al., “Nanopore electric snapshots of an RNA tertiary folding pathway,” Nat. Commun., vol. 8, no. 1, 2017.

[13] M. I. Rudenko et al., “Controlled gating and electrical detection of single 50S ribosomal subunits through a solid-state nanopore in a microfluidic chip,” Biosens. Bioelectron., vol. 29, no. 1, pp. 34–39, 2011.

[14] M. Rahman et al., “On demand delivery and analysis of single molecules on a programmable nanopore-optofluidic device,” Nat. Commun., vol. 10, no. 1, pp. 1–7, 2019.

[15] G. M. Cooper, “Translation of mRNA,” in The Cell: A Molecular Approach, 2nd ed., Sunderland (MA):Sinauer Associates, 2000.

[16] K. Bakowska-Zywicka and T. Twardowski, “Structure and function of the eukaryotic ribosome,” Cold Spring Harb Perspect Biol, vol. 4, p. a011536, 2012.

[17] J. R. Warner, P. M. Knopf, and A. Rich, “A Multiple Ribosomal Structure in Protein Synthesis,” Proc. Natl. Acad. Sci., vol. 49, no. 1, pp. 122 LP – 129, Jan. 1963.

[18] J. R. Warner and P. M. Knopf, “The discovery of polyribosomes,” Trends Biochem. Sci., vol. 27, no. 7, pp. 376–380, 2002.

[19] H. Noll, “The discovery of polyribosomes,” BioEssays, vol. 30, no. 11–12, pp. 1220–1234, 2008.

[20] E. E. Heyer and M. J. Moore, “Redefining the Translational Status of 80S Monosomes,” Cell, vol. 164, no. 4, pp. 757–769, 2016.

[21] A. C. Panda et al., “MiR-196b-mediated translation regulation of mouse insulin2 via the 5′UTR,” PLoS One, vol. 9, no. 7, pp. 1–11, 2014.

[22] A. C. Panda, J. L. Martindale, and M. Gorospe, “Polysome Fractionation to Analyze mRNA Distribution Profiles,” Bio Protoc, vol. 7, no. 3, p. e2126, 2017.

[23] H. Chassé, S. Boulben, V. Costache, P. Cormier, and J. Morales, “Analysis of translation using polysome profiling,” Nucleic Acids Res., vol. 45, no. 3, p. e15, 2017.

[24] I. T. Pereira et al., “Polysome profiling followed by rna-seq of cardiac differentiation stages in hescs,” Sci. Data, vol. 5, pp. 1–11, 2018.

[25] H. A. King and A. P. Gerber, “Translatome profiling: Methods for genome-scale analysis of mRNA translation,” Brief. Funct. Genomics, vol. 15, no. 1, pp. 22–31, 2016.

[26] C. Branco-Price, R. Kawaguchi, R. B. Ferreira, and J. Bailey-Serres, “Genome-wide analysis of transcript abundance and translation in arabidopsis seedlings subjected to oxygen deprivation,” Ann. Bot., vol. 96, no. 4, pp. 647–660, 2005.

[27] S. Liang et al., “Polysome-profiling in small tissue samples,” Nucleic Acids Res., vol. 46, no. 1, p. e3, 2018.

[28] R. J. Britten and R. B. Roberts, “High-resolution density gradient sedimentation analysis,” Science (80-.)., vol. 131, no. 3392, pp. 32–33, 1960.

[29] H. Yoshikawa et al., “Efficient analysis of mammalian polysomes in cells and tissues using ribo mega-SEC,” Elife, vol. 7, pp. 1–26, 2018.

[30] D. Simsek et al., “The Mammalian Ribo-interactome Reveals Ribosome Functional Diversity and Heterogeneity,” Cell, vol. 169, no. 6, pp. 1051-1065.e18, 2017.

[31] R. Aviner et al., “Proteomic analysis of polyribosomes identifies splicing factors as potential regulators of translation during mitosis,” Nucleic Acids Res., vol. 45, no. 10, pp. 5945–5957, 2017.

[32] M. Reschke et al., “Characterization and analysis of the composition and dynamics of the mammalian riboproteome,” Cell Rep, vol. 4, no. 6, pp. 1–7, 2013.

[33] E. Behrmann et al., “Structural snapshots of actively translating human ribosomes,” Cell, vol. 161, no. 4, pp. 845–857, 2015.

[34] D. Perry, D. Momotenko, R. A. Lazenby, M. Kang, and P. R. Unwin, “Characterization of Nanopipettes,” Anal. Chem., vol. 88, no. 10, pp. 5523–5530, May 2016.

[35] S. Zhang, M. Li, B. Su, and Y. Shao, “Fabrication and Use of Nanopipettes in Chemical Analysis,” Annu. Rev. Anal. Chem., vol. 11, no. 1, pp. 265–286, 2018.

[36] C. C. Chau, S. E. Radford, E. W. Hewitt, and P. Actis, “Macromolecular Crowding Enhances the Detection of DNA and Proteins by a Solid-State Nanopore,” Nano Lett., vol. 20, no. 7, pp. 5553–5561, 2020.

[37] S. W. Kowalczyk, D. B. Wells, A. Aksimentiev, and C. Dekker, “Slowing down DNA translocation through a nanopore in lithium chloride,” Nano Lett., vol. 12, no. 2, pp. 1038–1044, 2012.

