## Supplementary Information for "Ribosome fingerprinting with a solid-state nanopore"

Additional Figures

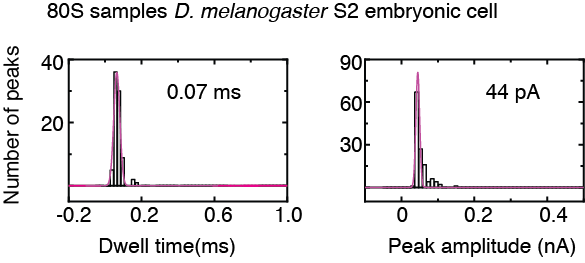

Figure S1 Dwell time and peak amplitude histogram for 80S ribosome samples from D. melanogaster S2 embryonic cell line.

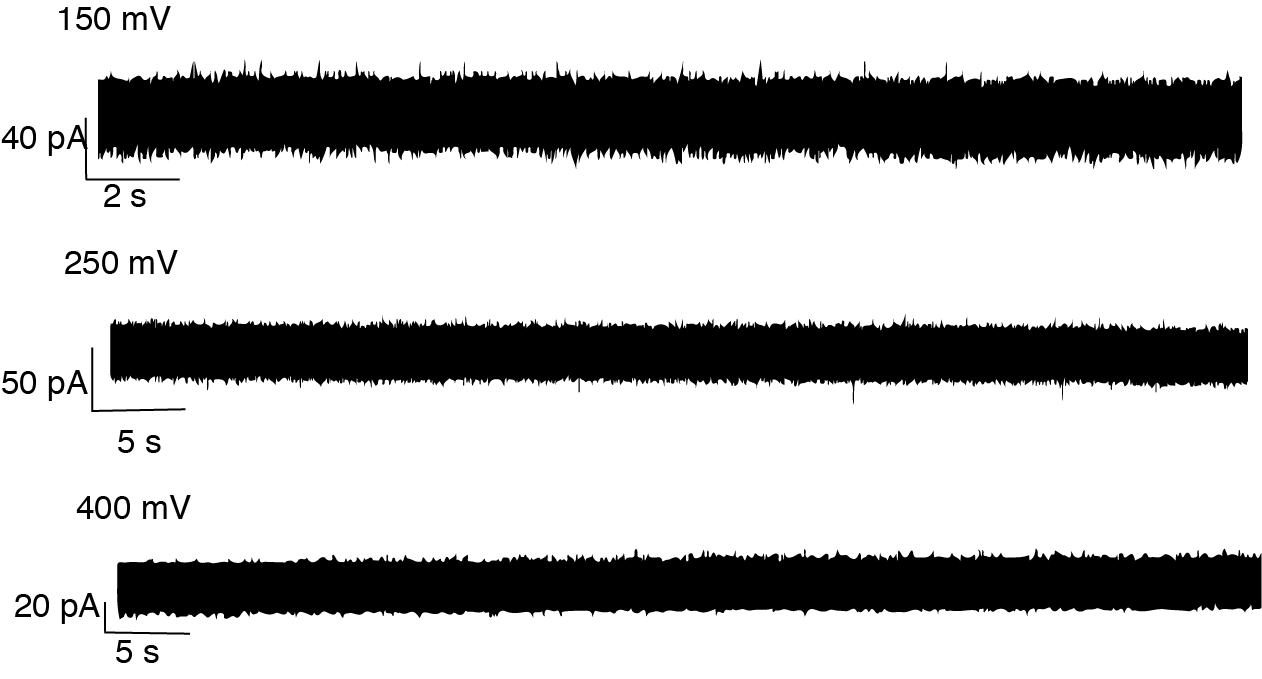

Figure S2 Ion current traces obtained at different voltages (150 mV, 250 mV and 400mV) with 40S samples from D. melanogaster S2 embryonic cell line. No events were observed.

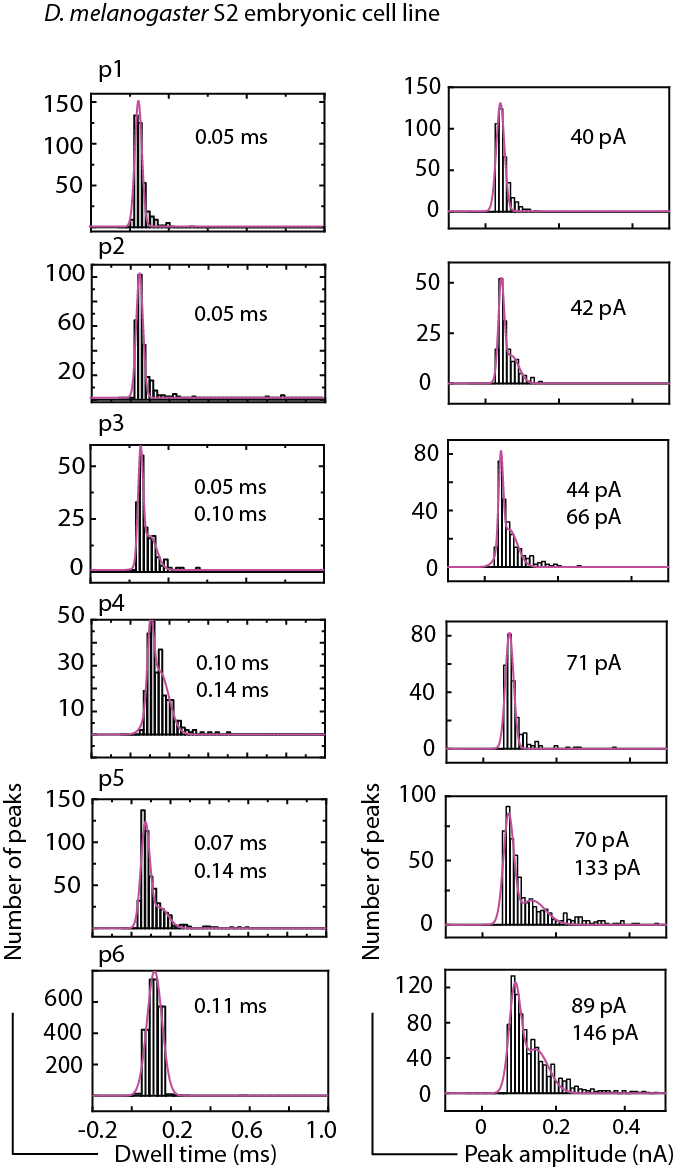

Figure S3 Individual histograms of dwell time and peak amplitude observed for the different polysome samples extracted from D. melanogaster S2 embryonic cells. The pink line follows the gaussian fit.

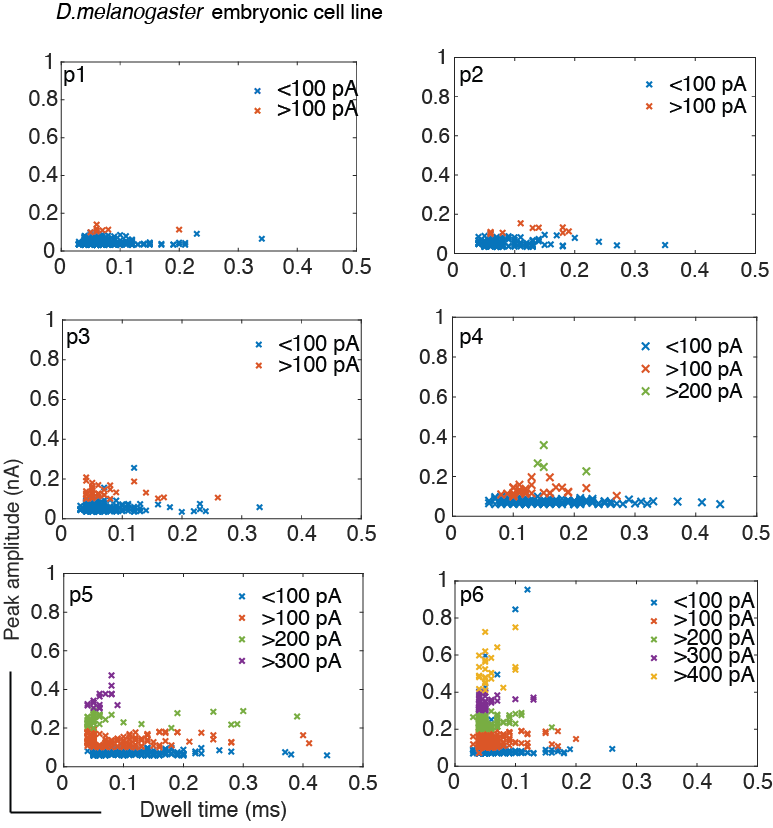

Figure S4 Scatter plots of peak amplitude plotted against dwell time for the D. melanogaster S2 cell samples. The events are colour-coded every 100 pA to highlight the increase in peak amplitudes.

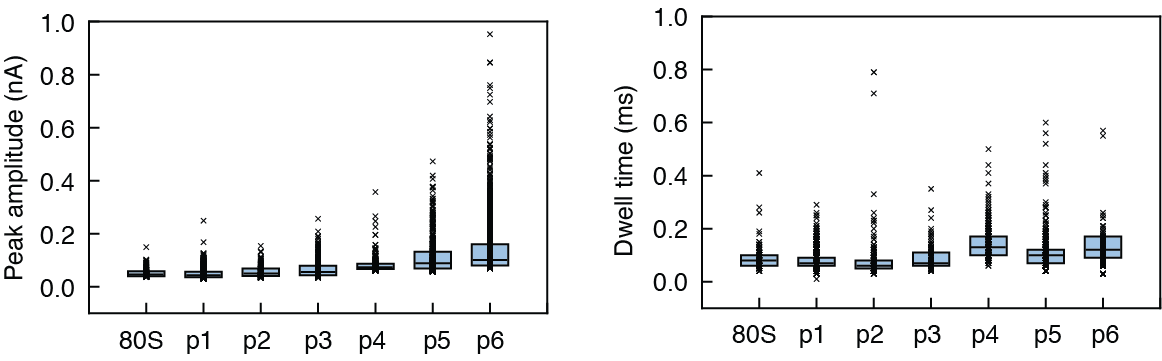

Figure S5 Peak amplitude and dwell time data for the 80S ribosome and various polysome samples of D. melanogaster S2 cells. The data represent >100 events for each sample. Error bars indicate standard deviation of the sample mean.

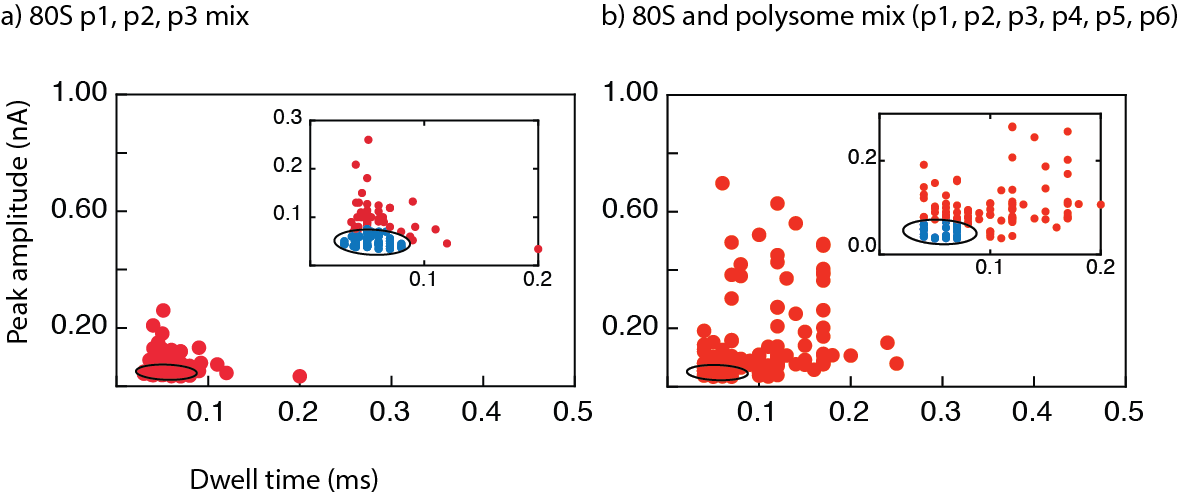

Figure S6 Scatter plots of peak amplitude plotted against dwell time for ribosome and polysome mixture from S2 cells, the data contains >100 events for each sample. The 95% confidence ellipse fitted for individual h80S data is used as boundary to differentiate h80s from the mixture.

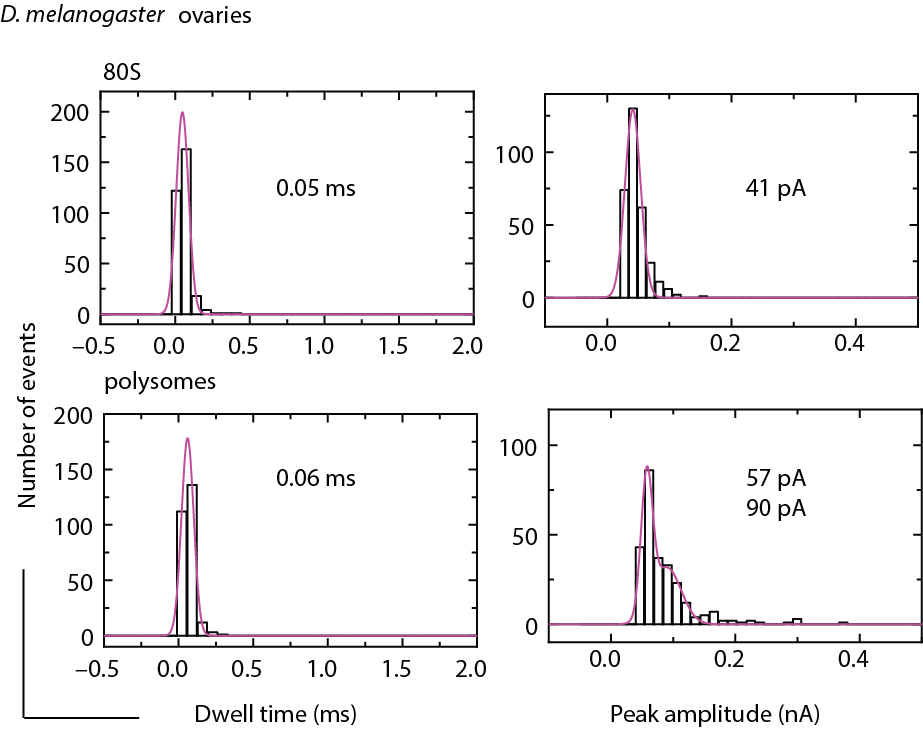

Figure S7 Histograms of peak amplitude and dwell times observed for 80S and polysome samples extracted from D. melanogaster ovaries. The pink line follows the gaussian fit.

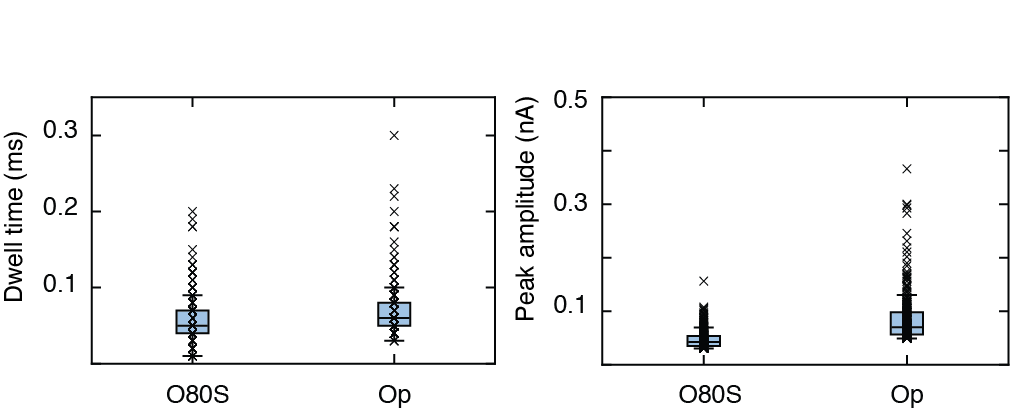

Figure S8 Individual dwell time and peak amplitude data plot for D. melanogaster ovaries samples (ribosome O80S and polysomes Op), the data represents >100 events from a 2 minute ion current trace for each sample. The error bars indicate the standard deviation of the sample mean.

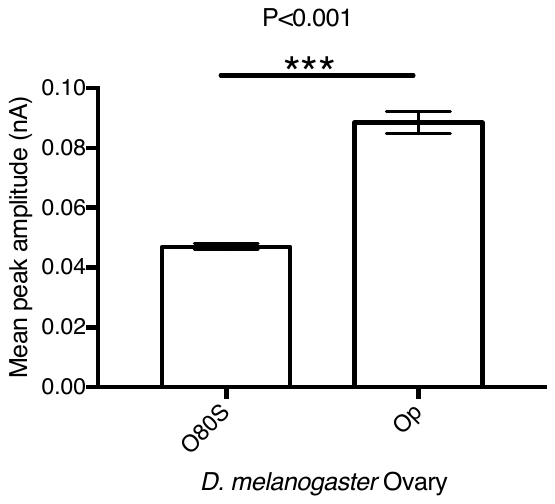

Figure S9 Mean peak amplitude data for D. melanogaster ovaries samples O80S and Op exhibiting a significant difference with p<0.001 represented with ***. Mann Whitney t-test was used for this data and the error bars represent standard error of mean.

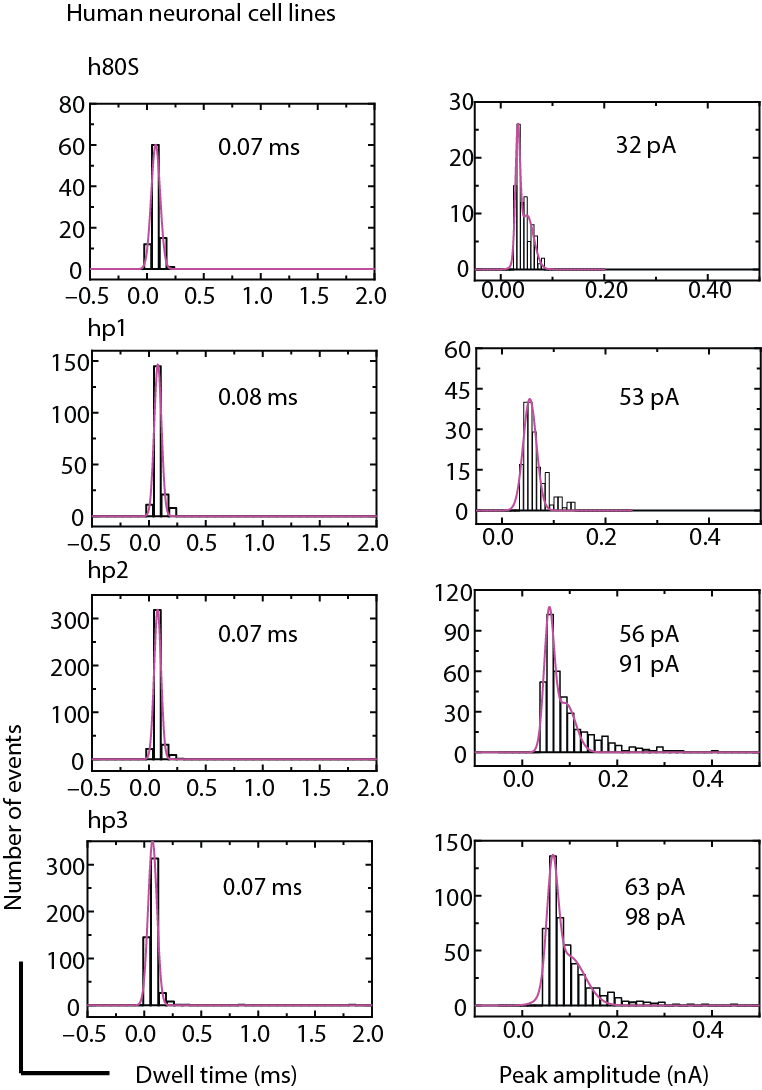

Figure S10 Individual dwell time and peak amplitude histogram data for single ribosome and polysome samples obtained from human human SH-SY5Y cells. The pink line follow the gaussian fit.

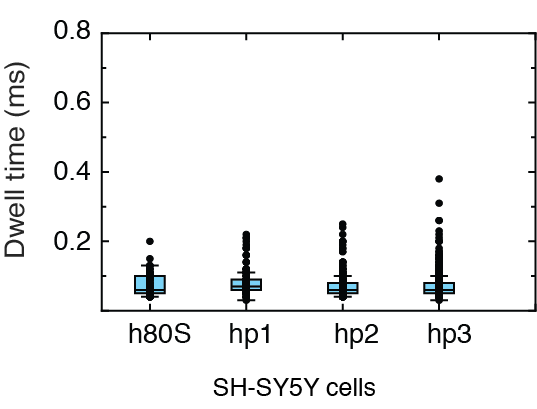

Figure S11 Dwell time data for 80S ribosome and polysome samples, the plot consists of >100 events for each sample.

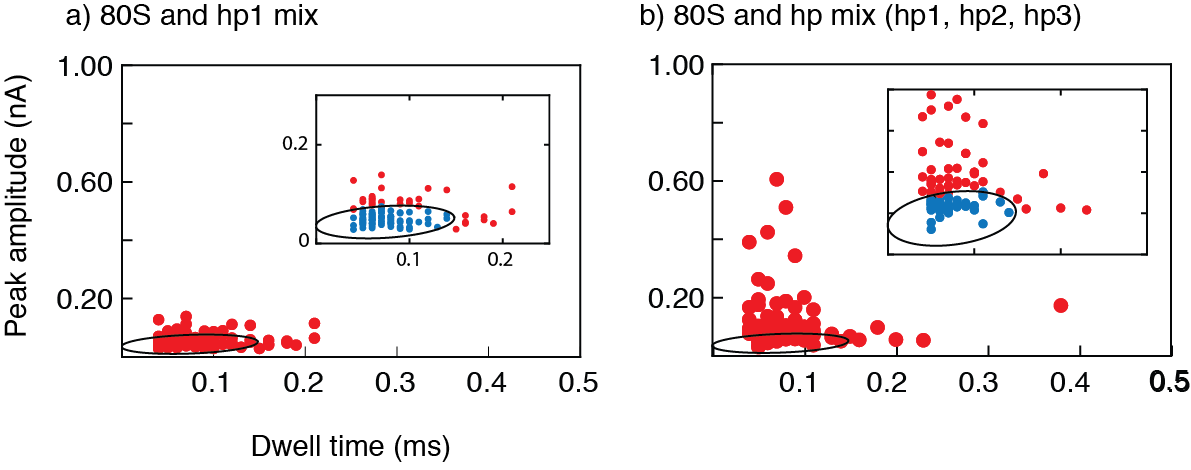

Figure S12 Scatter plots of peak amplitude plotted against dwell time for ribosome and polysome mixture from human SH-SY5Y cells, the data contains >100 events for each sample. The 95% confidence ellipse fitted for individual h80S data is used as boundary to differentiate them from the mixture.

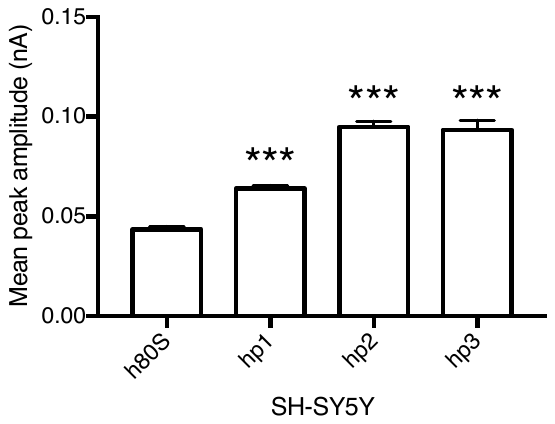

Figure S13 Human neuronal cell line samples. Mean peak amplitude data of SH-SY5Y polysomes compared with 80S samples exhibit a significant difference of p<0.001 represented by ***. Kruskal-Wallis test was performed for this data and the error bars indicate the standard error of mean.
